# Microbiome Stability with Chronic SIV Infection in AIDS-resistant Sooty Mangabeys

**DOI:** 10.1101/780825

**Authors:** RM Bochart, G Tharp, S Jean, AA Upadhyay, MM Crane, TH Vanderford, AM Ortiz, A Ericsen, JK Cohen, SE Bosinger

**Affiliations:** Division of Animal Resources, Yerkes National Primate Research Center; Yerkes NHP Genomics Core Laboratory, Yerkes National Primate Research Center; Division of Microbiology & Immunology, Yerkes National Primate Research Center; Barrier Immunity Section, Laboratory of Viral Diseases, Division of Intramural Research, NIAID, NIH; Department of Pathology & Laboratory Medicine, School of Medicine, Emory University

**Author notes:** To whom correspondence should be addressed: Dr. Steven E Bosinger, Yerkes National Primate Research Center, 954 Gatewood Rd, Atlanta, GA 30329, e-mail address.

## Abstract

Sooty mangabeys (SMs) are a natural host species of simian immunodeficiency virus (SIV) and avoid acquired immune deficiency syndrome (AIDS) despite persistently high viral loads, making them a pivotal research model for HIV pathogenesis. Unlike pathogenic SIV infection of macaque species, or HIV infection of humans, SIV-infected SMs maintain gastrointestinal barrier integrity. Here, we characterize the gastrointestinal bacterial microbiota of SIV-infected and uninfected SMs and perform a comparative analysis of diet-matched, rhesus macaques (RM). We assessed the fecal microbiome in fifty SM and thirty RM in total, and conducted analyses of the effect of SIV-status, species, and housing. When examining indoor-outdoor and indoor-only housing in our SM cohorts, biodiversity reduction and mild phylogenetic taxonomic perturbances were present. No statistically relevant differences were seen for biodiversity richness and evenness, or phylogenetic taxonomic communities between SIV negative and positive SM cohorts. In contrast, with pathogenic early chronic SIV infections in RM a trend of alpha diversity loss and increase of beta diversity and few phyla taxonomic communities differed. Lastly, we observed lower levels of pathobiont bacterial communities in SIV-uninfected SMs relative to RMs. These data suggest that the pre-existing bacterial community structure may contribute to the divergent phenotype between SIV natural hosts and pathogenic macaque species.

**Importance:** Human immunodeficiency virus remains a global concern. The sooty mangabey (SM) monkey is an important biomedical research model for understanding HIV pathogenesis due to its ability to avoid AIDS disease progression despite high viremia. In people living with HIV, gastrointestinal dysbiosis towards enrichment of pathobiont communities has been frequently reported. In this study we characterized the fecal microbiota of a primate non-pathogenic SIV host, the SM, and made direct comparisons to a pathogenic SIV host species, the rhesus macaque. We observed that SMs exhibit stability of the microbiota community into chronic SIV infection, which contrasts with SIV-infected rhesus macaques, in which we observed bacterial community divergence relative to uninfected animals. Collectively, our observation of stabilization of beneficent taxa in the mucosa of AIDS-resistant primates suggests that therapeutic strategies to enrich these communities may have potential for ameliorating the gastrointestinal inflammation in people living with HIV.

## Introduction

Chronic immune activation (IA) has long been considered to be a primary factor driving disease in HIV infection(1, 2). IA is a barrier to achieving optimal health for people living with HIV (PLWH) on suppressive antiretroviral therapy (ART)(3, 4). Residual inflammation remaining after ART is predictive of several non-AIDS pathogenic sequelae such cardiovascular disease, incomplete CD4 T cell reconstitution, and premature immune senescence(3, 4). Despite the influential role that chronic IA exerts on disease in untreated and ART-controlled HIV infection, its underlying causes remain largely uncharacterized.

Natural host NHP species are African primates that remain AIDS-free despite life-long SIV infection and have been used for many years as a model to study HIV-related inflammation(5). The African sooty mangabey (SM) monkey (*Cercocebus atys*) is one of the best studied natural hosts of simian immunodeficiency virus (SIV). The impact of SIV infection on the gastrointestinal mucosa differs significantly between natural hosts and Asian macaques, and is likely a critical site underlying the divergent response to SIV between species (6, 7). In pathogenic SIV infections, mucosal CD4+ T cells are rapidly depleted during acute infection, and remain at levels significantly lower than baseline(8, 9). In contrast, natural SIV hosts stabilize, or even partially recover, mucosal CD4+ T cells(6, 7). Observable gut pathology is virtual absent in natural hosts after SIV infection – but is highly prevalent in HIV infection and SIV infection of Asian macaques(10). Over the past decade, a well-accepted model has emerged to explain the chronic, persistent immune activation in pathogenic HIV/SIV infection: inflammation occurring in acute infection compromises the integrity of the intestinal barrier and allows a perpetual influx of immunostimulatory microbial content from the intestinal lumen to the systemic circulation(11, 12). Microbial translocation, immune activation and barrier damage cells do not normalize in humans or NHPs even after long-term suppressive antiretroviral therapy(4). Natural host species, however, do not exhibit mucosal pathology, epithelial barrier breakdown, or persistent infiltration of microbial products into the lamina propria, although transient levels of microbial products in the plasma are observed in acute SIV infection(6, 7). The mechanisms by which natural hosts are able to preserve mucosal integrity have not been delineated. One putative mechanism is based on an observation that natural hosts preferentially retain of mucosal CD4+ TH17 and innate lymphoid cells, which promote epithelial barrier structural integrity(13). Recently, we described a conserved polymorphism in the innate immune receptor TLR4 in multiple natural host species, which dampens signaling to LPS and other TLR4 ligands, which could conceivably contribute to decreased gastrointestinal inflammatory response after SIV infection(14).

The profound disruption of mucosal immunity by HIV/SIV infection has instigated several studies on its impact on the mucosal microbiome in humans and NHPs(reviewed in (15, 16)). In PLWH, a general reduction of gastrointestinal microbiota richness, bacterial community dysbiosis, and opportunistic pathogenic taxa enrichment relative to healthy controls has been consistently reported(17-19). HIV patients on ART have been observed to retain a persistent perturbance of microbiota and enrichment of potentially pathogenic bacteria(17, 18), which has been linked to immunological factors potentiating mucosal inflammation(18). Mucosal dysbiosis has also been reported with SIV infection of non-human primates, however it is not consistently observed(20-24). In the setting of ART, SIV-infected NHPs have been reported to have microbial dysbiosis that reverts towards the pre-SIV microbiota state over time(20).

The goal of this study was to test the hypothesis that the mucosal microbiome of the SM would be different from non-natural hosts, and potentially contribute to the AIDS-resistant “natural-host” phenotype. Specifically, we tested if the lack of inflammation and gut pathology in SIV-infected SMs was due to alterations of the gut microbiome that enriched non-inflammatory/commensal bacterial species relative to non-natural host rhesus macaques. We also tested the impact of indoor vs. indoor-outdoor housing on the SM microbiome in a diet-controlled setting. To address these questions, we assessed the microbiome associated with stool collected from 80 captive NHPs (50 SMs and 30 RMs). Our results demonstrate that, relative to RMs, the microbiome in SMs remains stable during SIV infection. Moreover, the microbiome in uninfected SMs is enriched in beneficent taxa, and harbours lower levels of pathobiont taxa relative to uninfected RMs. These data support a hypothesis in which the microbiome may contribute to an attenuated inflammatory environment in the mucosa of AIDS-resistant natural host species.

## Results

### Animal Cohorts

An overview of the study design is depicted in **Figure 1A** and summarized in **Table 1**. Our study included a total of fifty SMs to characterize the gastrointestinal microbiome of an SIV natural host species. Specifically, we tested the impact on fecal microbiota by (i) SIV status, and (ii) captivity lifestyles (outdoor vs indoor) within a diet-controlled research setting. We enrolled thirty gender balanced SMs living in indoor-outdoor social housing with equal matched groups of SIVsmm-infected (n = 15) and uninfected controls (n = 15). Additionally, we included twenty SMs living in indoor-only standard cage housing with 15 SIVsmm-infected (10 males/5 females) and five healthy controls (2 males/3 females). All SIVsmm-infected SMs were in chronic phase of infection (28 animals >10 years post-infection, with two animals with an undetermined d.o.i. but confirmed >3 yrs.). To control for genotypes known to influence SIV infection, we excluded animals that were homozygous for the CCR5Δ24 mutation that abrogates CCR5 surface expression(25) (25 wt/wt, 0 wt/Δ24) and animals we had genotyped as homozygous for a 2-nt deletion (CCR5Δ2) (23 wt/Δ2). Two SMs were of unknown CCR5 genotype. For the purposes of comparing the microbiome in natural vs. non-natural NHP host species, we obtained fecal samples from a gender-balanced cohort of 15 Indian RMs with equal matched groups of SIVmac239-infected (38-51 days post infection, n = 15) and uninfected control RMs (n = 15). All NHPs were maintained on a matched, diet controlled environment and were greater than six months from of administration of antibiotic therapy. The SIVmac239 infected RM and outdoor-indoor SIVsmm SM cohorts were ART naïve, whereas 10/15 of the indoor-only cohort of SIVsmm SMs had been administered ART as part of a prior study(26), but were at least 2.5 years post ART administration at the time of sample collection. A detailed list of animals used in this study and relevant metadata is included in Supplementary Table 1.

**Table 1.** Summary data of study animals.

| Cohort | *n* | Sex | Age (years) | Mean<br>Plasma<br>Viral Load<br>(RNA<br>copies/mL) | Duration<br>SIV+ |
| --- | --- | --- | --- | --- | --- |
| SM<br>SIV negative | 20 | M (n=9)<br>F (n=11) | Median: 18<br>Min: 13<br>Max: 24 | N/A | N/A |
| SM<br>SIV positive | 30 | M (n=18)<br>F (n=12) | Median: 21<br>Min: 5<br>Max: 35 | $1.2 \times 10^5$ <sup>a</sup> | >10 yrs (n=28)<br>>7 yrs (n=1)<br>>3 yrs (n=1) |
| RM<br>SIV negative | 15 | M (n=7)<br>F (n=8) | Median: 6<br>Min: 4<br>Max: 7 | N/A | N/A |
| RM<br>SIV positive | 15 | M (n=7)<br>F (n=8) | Median: 3<br>Min: 3<br>Max: 4 | $1.9 \times 10^6$ | 38-51 days |

<sup>a</sup> – plasma viral load data was available for 22/30 animals.

**Fig 1.**
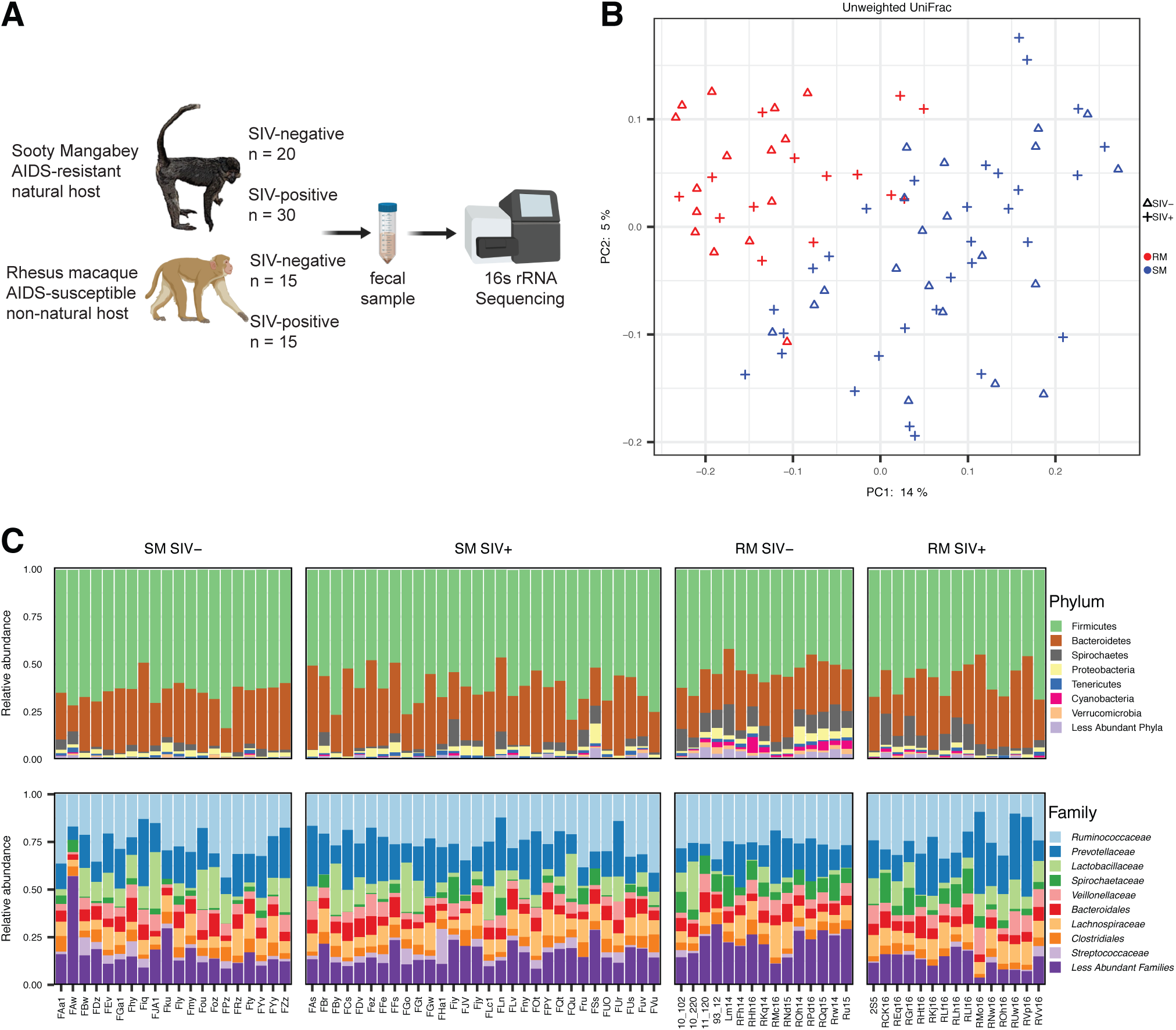
Gastrointestinal microbiota stability with chronic SIV infection in the natural SIV host. **A)** Study design cartoon depiction; includes species numbers, SIV infection status of study animals, and overview on fecal sample processing. **B)** Principal coordinate plot of Unweighted UniFrac distances (a measurement of beta diversity) comparing SMs (blue) with RMs (red) and SIV status (negative= Δ, positive=+). **C)** Bar graph representation of phylogenetic phyla and family bacterial community acquisition. Each individual animal represents a single bar and grouped according to species (SM on left, RM on right) and SIV status (SIV-, SIV+).

### Microbiome biodiversity reduction and perturbance is seen with increased captivity lifestyles within a research setting

Due to the limited availability of SMs in the uninfected control cohort within our indoor facility (n=5), cohorts of SMs were assembled from two sites within the Yerkes National Primate Research Center (YNPRC), the breeding facility, where animals live in indoor-outdoor enclosures with access to natural substrates, and our primary research location, where they are housed in standardized indoor-only cage housing. Previous studies have shown a reduction in alpha diversity within the fecal microbiome of captive *Cercopithecidae* species compared to those in the wild(27, 28), thus we first investigated if caging influenced the microbial composition of the SMs. We examined for potential microbiota differences between captivity lifestyles within a research setting by comparing twenty indoor-only singly housed SM, and thirty socially housed outdoor-indoor SMs. When comparing the indoor vs outdoor irrespective of SIV status, we observed that the overall composition of the fecal microbiota was highly similar for the most predominant phyla, Firmicutes and Bacteroidetes (**Fig. S1A**). Firmicutes accounted for 64% and 61% of the sequenced taxa, in the outdoor-indoor and indoor samples, respectively, and Bacteroidetes taxa were estimated at 30% and 28%, although these differences were not statistically significant (**Fig. S1C**). At the family level, the abundant taxa within the indoor and outdoor-indoor groups were *Ruminococcaceae (*26% vs 26%, ns*), Prevotellaceae* (17% vs 19%, ns), *Lachnospiracheae* (9% vs 8%, ns), *Lactobacillaceae* (5.5 % vs 11.8%, p-value=0.001). We observed one animal (FAw) with a highly diverse, polymicrobial distribution of taxa, but this was atypical. Indoor-only housing trended towards reduced alpha diversity of gastrointestinal bacterial community assessed by both the chao1 (richness) and Shannon (richness and evenness) indices (**Figure S1B**). Despite the similarity of the predominant taxa, when we compared SM cohorts by lifestyle using the Bray-Curtis metric (a measurement for dissimilarities between study cohorts) for beta diversity we observed overt microbial community differences (**Fig. S1E**). To examine for the additional taxa underlying the divergence of beta diversity, we tested for differences in taxa at the phyla level between lifestyles and observed modest but significant reductions in Spirochaetes and Proteobacteria in the outdoor animals (**Fig. S1C**). At the genera level, we observed that indoor housed SMs had a significant reduction *Lactobacillus* and *Streptococcus* (**Fig. S1D**). Taken together, these data indicate that a mild microbial biodiversity loss and minor taxonomic compositional shift was associated with indoor-only housing. Based on this observation of minimal impact of housing on diversity and community structure, we were able to pool SMs from both housing groups into cohorts based on SIV-infection status for subsequent analyses.

### The fecal microbiome in natural host SMs remains stable in chronic SIV infection

Based on the observation of minimal impact of housing on diversity and community structure, we pooled data from SMs of both housing groups for subsequent analyses based on SIV-infection status (**Fig. S1A-E, Fig. 2A**). One of the key physiological differences between primates that develop AIDS (humans, *Macaca* species) and SIV natural hosts, is that the latter maintains a healthy intestinal barrier despite the loss of mucosal CD4+ T cell subsets. We hypothesized that SIV-infected SMs would not exhibit the microbial dysbiosis reported in some studies for macaque species(29). At the phyla levels, the community structure between uninfected and SIV-infected SMs was largely similar and there was no significant differences between the levels of the predominant taxa Firmicutes (64% vs 61.5%, ns) or Bacteroidetes (28.7% vs 29.9, ns), nor differences in the next two most prevalent phyla, Spirochaetes (2.5% vs 3%) and Proteobacteria (2.1% vs 2.5%) (**Fig. 1C, Fig. 2B**). The composition of microbiota was also stable at the family level. The most abundant taxa were not observed at significantly different levels between SIV- and SIV+ cohorts: *Ruminococcocaceae* (26.4% vs 25.4%, ns), *Prevotellaceae* (17.4% vs 18.8%, ns), *Lactobacillaceae* (10.2% vs 9%, ns) (**Fig. 1C**). We also examined the impact of SIV infection on overall diversity of the fecal microbiome. When comparing uninfected controls and chronically SIV-infected SMs, we observed no statistically significant differences in microbiota alpha diversity when examining operational taxonomic units (OTUs) as assessed by either the chao1 and Shannon metrics (**Fig. 2A**), as well as for beta diversity using the unweighted UniFrac metric which measures for differences between the cohorts (**Fig. 1B**). Similarly, we did not observe a difference in diversity metrics associated with SIV status when we compared the indoor and the outdoor-indoor groups separately **(Fig. 2A)**.

**Fig 2.**
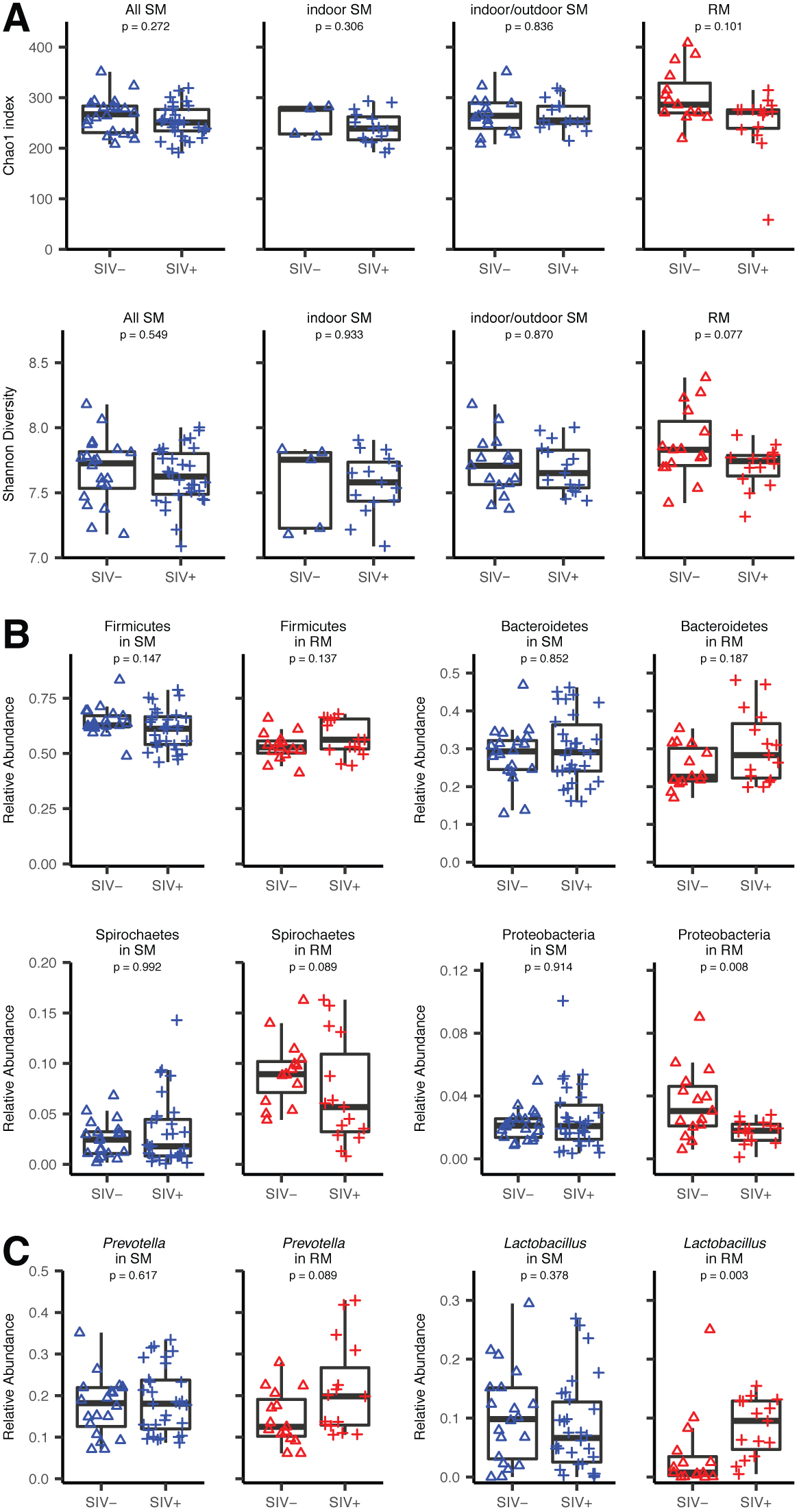
Modest microbiota biodiversity and phylogenetic bacterial community disruption with early chronic SIV infection in the non-natural SIV host, and microbiota stability in the natural host species. **A)** Chao1 index (a total OTU richness measurement for alpha diversity) was used to compare SIV negative (Δ) and SIV positive (+) SM (blue) cohorts separated by lifestyle (combination (left), indoor-only (middle-right), indoor-outdoor (middle-left)), and the RM (red) cohort (right), all data is compared by the paired Wilcoxon rank sum test. Shannon diversity indices (an alpha diversity metric) were used to measure for differences within the species cohorts in comparison to SIV status. The plots show SM cohorts separated by lifestyle and combined due to insignificant differences (All SM), and RM cohort (right). **B)** Relative abundance of the predominant phyla Firmicutes, Bacteroidetes, Spirochaetes, and Proteobacteria represented by box plots comparing SIV negative (Δ) and SIV positive (+) SM (blue) and RM (red) cohorts compared by the paired Wilcoxon rank sum test. **C)** Relative abundance of genera Lactobacillus and Prevotella represented by box plots comparing SIV negative (Δ) and SIV positive (+) the SM (blue) and RM (red) cohorts, which are compared by the paired Wilcoxon rank sum test.

We then assessed for global taxonomic genera differences and included a comparative analysis for each part of the study for *Lactobacillus* and *Prevotella*, as previous literature have reported these genera have a significant impact on mucosal immunity in SIV or HIV pathogenesis (29). The genus *Lactobacillus* has been correlated with the inhibition of IDO1 activity, and helping preserve mucosal TH17 CD4+ T cells and gastrointestinal mucosal integrity (30). Fecal *Lactobacillus* levels have been reported to decrease in pathogenic SIV infection of RMs, but increase in non-pathogenic infections of African green monkeys as they progress from acute to early stages of chronic infection (30). Here, when we contrasted SIV-negative vs SIV-infected SMs, we did not observe a significant difference in the levels of *Lactobacillus* (**Fig 2C**). We also tested for differences in the genus *Prevotella*, as it has been reported to be enriched in PLWH, and has been associated with chronic immune activation and inflammation (31). Here, we did not observe significant differences for *Prevotella* levels between chronically SIV infected SMs and uninfected animals (**Fig 2C**). Collectively, these data show that chronic SIV infection in SMs has a minimal impact on microbial diversity, bacterial community richness, and overall community structure. Similarly, we did not observe enrichment or loss of potentially beneficial or pathogenic taxa.

### Early chronic SIV infection in macaques is associated with lowered community diversity

A primary goal of our study was to test if the fecal microbiome of non-pathogenic SMs either harbours lower levels of pathobionts, or conversely, causes enrichment of beneficial taxa relative to pathogenic species. In order to make an effective interspecies comparison, we collected fecal samples from 30 diet-matched animals, 15 uninfected RMs and 15 RMs at the early stages of chronic SIV infection (38–51 d.p.i.). Overall, the community structure of the fecal microbiome was highly similar to that reported previously for rhesus (30), pig-tailed (20), and cynomolgus macaques (32), with the predominant phyla being Firmicutes, Bacteroidetes, Proteobacteria, and Spirochaetes (**Fig. 1C**), and was also similar to that described for SMs (**Fig. 1C**). Previous studies have described microbial dysbiosis, and reduced alpha diversity associated with SIV infection (22, 30). Similarly, we observed a trend of alpha diversity reduction, although at insignificant levels, represented by the chao1 and Shannon diversity metrics (**Fig. 2A**), as well as an increase of dissimilarities between the cohorts shown by the unweighted UniFrac metric (**Fig. 1B**). In contrast to prior reports(30), the phylogenetic phylum Proteobacteria’s abundancy significantly reduced with SIV infection (**Fig. 2B**), while all other phyla had no significant changes(**Fig. 2B**). At the genera level RMs show an insignificant increase of *Prevotella* and a significant increase of *Lactobacillus* with SIV infection (**Fig. 2C**). Overall, while we noted differences in the changes of some low abundance taxa associated with SIV infection, the major taxa we observed was largely concordant with prior reports of the macaque microbiome at different facilities. Based on this, we judged our RM dataset suitable for meaningful comparison with the SM microbiome.

### Diversity of the microbiome is higher in uninfected rhesus macaques relative to sooty mangabeys

Next, we directly compared the gastrointestinal microbiota of SMs with RMs to determine if natural host species have a reduction or enrichment of microbiota taxa that may underlie their mucosal immunological resistance to SIV progression. During acute SIV infection, both RMs and SMs exhibit mucosal CD4+ T cell loss, however SMs maintain integrity of the mucosal epithelial barrier(6). Assessment of the transcriptomic profile of mucosal tissue during very early SIV infection has demonstrated that RMs are characterized by inflammation, whereas natural hosts exhibit gene expression signatures consistent with wound healing (Barrenas *et al*. Nat. Commun. *in press*). Here, we tested if the microbiome in SMs and RMs, prior to infection, is different and could potentially contribute to this differential phenotype. We compared the RM (n=15) and SM (n=20) healthy control cohorts to examine for species differences. Bacterial species richness of OTUs were statistically significantly higher in RMs in comparison to SMs represented by the chao1 metric plot (p=0.042), and trended higher when assessed by the Shannon index (p=0.055) (**Fig. 3A**); this elevated richness in RMs is distinctly significant considering that the majority of SMs in this cohort exhibited an outdoor-indoor lifestyle (n=15/20). This diversity was reflected in the overall community structure, as SMs had a significantly higher proportion of Firmicutes phyla (64.2% vs 53.4%, p <0.001). In SMs, the two predominant phyla, Firmicutes and Bacteroidetes, accounted for 92.9% of taxa. Comparatively, uninfected RMs had a much more polymicrobial phylogenetic distribution (**Fig. 1C**), as only 78% of taxa were comprised of Firmicutes and Bacteroidetes, and 9/15 RMs had greater than 20% of their community phyla represented by Spirochaetes, Proteobacteria, and other less abundant microbes. The species cohorts showed distinct beta diversity differences with increased dissimilarity represented by the Bray-Curtis metric (**Fig. 3B**). Interestingly, SMs had lower levels of Spirochaetes (p<0.001) and Proteobacteria (p=0.059) relative to RMs (**Fig. 3C**). SMs samples had significantly higher abundance of *Lactobacillus* than RMs, but no significant difference for *Prevotella* abundance was seen between the species (**Fig. 3D**).

**Fig 3.**
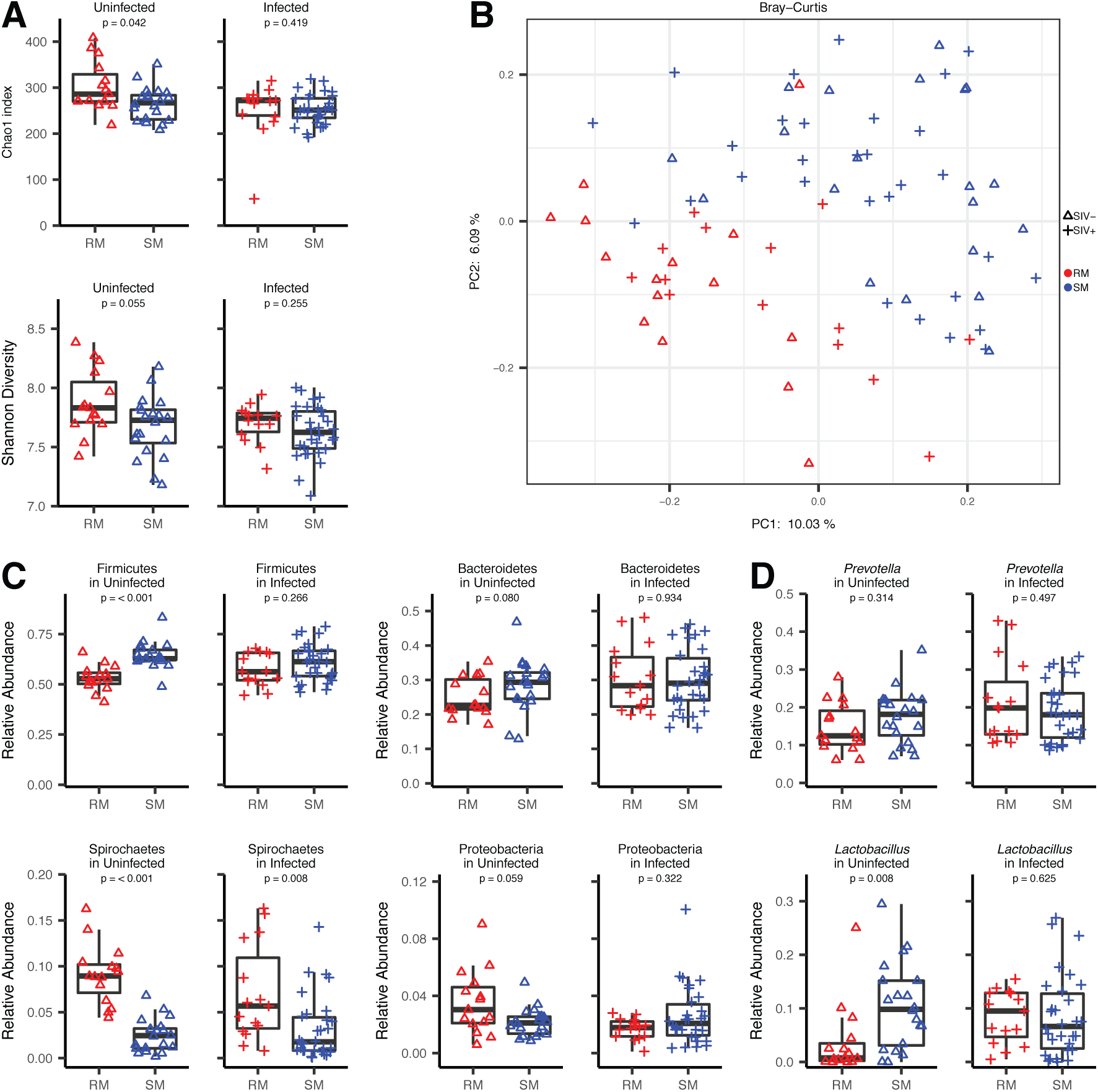
The fecal microbiome of uninfected RMs is enriched for pathobiont bacterial communities relative to SIV negative natural host species. **A)** Chao1 index (a total OTU richness measurement for alpha diversity) was used to compare SIV negative (Δ) RMs (red) and SMs (blue) (left figure), and SIV positive (+) RMs (red) and SMs (blue) (right figure). Shannon diversity (an alpha diversity metric) were used to compare SIV negative (Δ) RMs (red) and SMs (blue) (left figure), and comparing SIV positive (+) RMs (red) and SMs (blue) (right figure). Both alpha diversity measurements were compared by the paired Wilcoxon rank sum test. **B)** Principal coordinate plot of Bray-Curtis dissimilarities measurement of beta diversity comparing SIV negative (Δ) RMs (red) and SMs (blue), and comparing SIV positive (+) RMs (red) and SMs (blue). **C)** Relative abundance of the predominant phyla Firmicutes, Bacteroidetes, Spirochaetes, Proteobacteria represented by box plots comparing SIV negative (Δ) or SIV positive (+) RMs (red) and SM (blue) cohorts compared by the paired Wilcoxon rank sum test. D) Relative abundance of genera Lactobacillus, Prevotella, Streptococcus represented by box plots comparing SIV negative (Δ) or SIV positive RMs (red) and SMs (blue) cohorts compared by the paired Wilcoxon rank sum test.

### The microbiome of RMs and SMs during chronic SIV infection is similar due to loss of taxa in RMs

One of the hallmarks of lentiviral infection of natural host species is the lack of systemic immune activation that is seen in pathogenic HIV/SIV infection. Here, we tested the hypothesis that the natural host SMs maintain lower levels of microbial taxa associated with inflammatory responses by direct comparison of the community composition with RMs during chronic infection. We contrasted the fecal 16s rRNA profiles of 30 chronically SIV-infected SMs (>10 years (n=28), >3 years (n=2)) SIV-infected SMs with 15 SIVmac239 infected RMs (38-51 d.p.i.). Unlike the comparison of uninfected SMs with RMs, when comparing samples from infected animals, we observed no major differences in the most abundant Firmicutes and Bacteroidetes phyla, nor any significant differences at the family level (**Fig. 1A, 3C**). Likewise, there were no significant differences in alpha diversity between SIV-infected SMs and RMs (**Fig. 3A**). The cohorts were not completely indistinguishable, as demonstrated by Bray-Curtis dissimilarity metric (**Fig. 3C**), however the differences were more modest than contrasts of uninfected SMs and RMs. The lack of a difference between SMs and RMs was due to a loss of taxa diversity in the SIV-infected RMs relative to uninfected animals. When we compared individual taxa for differences, the levels of Proteobacteria were found to be lower in RMs than in SMs, this result was the opposite trend of that observed for uninfected animals (**Fig. 3C**). Unlike most of the less abundant taxa, Spirochaetes levels were retained in most RMs, and this taxa remained significantly higher compared to SMs (**Fig. 3C**). Based on reports of the importance of *Lactobacillus* and *Prevotella* to HIV infection, we tested for differences between species, but observed no significant differences (**Fig. 3D**).

## Discussion

This study represents the first comprehensive characterization of the gastrointestinal microbiome in captive natural SIV host species sooty mangabeys (SM). We observed that chronic SIV infection in SMs is associated with gastrointestinal microbiota stability of bacterial community richness, biodiversity, and phylogenetic taxa. In contrast, in early chronic pathogenic SIV infection of RMs, we observed a biodiversity difference with a decreased alpha diversity, a trend to increased beta diversity, and taxonomic phyla community perturbances, most importantly Proteobacteria depletion and *Lactobacillus* enrichment. Further, this study includes interspecies comparison of uninfected healthy RMs and SMs, with RMs exhibiting microbiota diversity expansion, overt beta diversity dissimilarities between the species, a phylogenetic taxa higher yield of Proteobacteria and Spirochaetes, and reduction of Firmicutes and *Lactobacillus* in comparison to the SM. When performing interspecies comparison of infected animals, beta diversity continues to have dissimilarities (although lessened) between the cohorts, with RMs showing expansion of phyla Spirochaetes, other differences reported for uninfected species comparison do not remain significant. Lastly, we conducted a retrospective analysis of the impact of caging strategy and lifestyle, and observed that SMs living within indoor-only standard housing had alpha diversity reduction, increased beta diversity, and microbial community expansion of Proteobacteria and Spirochaetes, and reduction of genera *Streptococcus* and *Lactobacillus* compared to the outdoor-indoor SM cohort. Although SMs did not show significant enrichment of immunological beneficial bacteria after SIV infection, we did observe enrichment of beneficial bacterial communities and reduction of pathobiont populations in SIV-negative SMs relative to non-natural host RMs. Consistent with our second hypothesis, we did discover intraspecies microbiome changes between lifestyle differences within a captive research setting in SMs.

Microbiota dysbiosis typically manifests as a perturbance of commensal bacteria towards pathobiont communities and a loss of microbiota diversity(33). A reduced gastrointestinal microbiota alpha diversity has been linked to mucosal, metabolic, cognitive, endocrine, and immune diseases (33-36). A decrease in alpha diversity(37-39) and an increase in beta diversity has been reported in several studies of untreated HIV patients compared to healthy controls(38, 39). Similarly, several studies have reported a loss of alpha diversity in SIV infection of macaques (30) or HIV in humans (37, 38, 40), and here, we have also observed a trend to reduced diversity in pathogenic SIV infection. In contrast, chronic infection of natural hosts was not associated with a loss of diversity or perturbation of community structure, rather, we observed a stability of microbiota alpha and beta diversity and lack of dysbiosis. Some evidence has suggested that microbiome stability may be associated with lowered pathogenesis in humans: as antiretroviral therapy-naïve, HIV-infected, elite controller-patients (i.e. with undetectable plasma viral loads) have been shown to have gastrointestinal microbiota diversity resembling that of healthy controls(37, 38). Metagenomic sequencing of AGMs during acute SIV infection demonstrated no changes associated with the gastrointestinal virome or bacterial microbiome(24). Vujkovic-Cvijin *et al* reported that fecal *Lactobacillus* abundancy was enriched in early chronic SIV infection of AGMs (defined as 46-56 days) (30), which we did not see this in our data, although the majority of the SMs in our study had been SIV-infected for >10 years. Overall, our study demonstrated that SMs with chronic SIV infection were observed to have stability of the gastrointestinal bacterial community, in contrast to diet-matched RMs, which exhibited a significant fluctuation of several taxa after SIV infection. While our data does not address mechanism, in the context of previous literature findings, it does support a model in which microbiome stability is linked to low immune activation in HIV/SIV infection.

Several studies have investigated the impact of SIV infection on the fecal microbiome in macaque species, and while some of these findings have been inconsistent with one another in terms of observed dysbiosis or changes in specific taxa, likely due to differences in species (rhesus, cynomolgus, or pig-tailed have been reported), timing of collection, or technical variables(20-24), translocation of Proteobacteria has been consistently reported (20, 32). Here, we observed that: (i) levels of Proteobacteria in uninfected RMs trended higher than in uninfected SMs; (ii) compared to uninfected RMs, macaques sampled during early chronic SIVmac239 infection (38-51) have lower Proteobacteria; (iii) Proteobacteria levels in SMs were not significantly changed between infected and uninfected animals. Our observation of lowered Proteobacteria levels in RMs during early chronic infection is consistent with the findings of Klase *et al*, who reported depletion of colonic Proteobacteria levels at ∼40 days d.p.i., that subsequently normalizes, although incompletely, when sampled at day 90. Proteobacteria are comprised of several pathogenic, Gram-negative taxa and have been linked to multiple pro-inflammatory gastrointestinal disorders including Crohn’s disease and inflammatory bowel disease (41). In the context of SIV infection, lymphoid tissue levels of Proteobacteria correlate with immune activation (assessed by HLA-DR+CD4+ T cells) in SIV infected macaques(20). In recent work, we have observed that during hyperacute SIV infection (< 3 d.p.i.) RMs have a transcriptional signature in the mucosa consistent with TLR4-driven inflammation that is absent in natural host AGMs. We have also identified a mutation in TLR4 in SMs that is conserved in multiple natural host species(14). In this regard, it is intriguing that we observed significantly lower levels of Proteobacteria in uninfected SMs compared to RMs, and supports a hypothesis in which SMs harbour a gastrointestinal microbiome less prone to supporting inflammation. Moreover, our observation that Proteobacteria remains largely unchanged between chronically SIV-infected and uninfected animals reinforces the observation that natural host species maintain an intact gut barrier.

Several beneficial characteristics have been attributed to *Lactobacillus* spp. including anti-inflammatory and immunoprotective properties, reducing neutrophil life span, and inhibition of local pathobiont overgrowth (42, 43). A reduction of *Lactobacillus* with SIV or HIV infection has been reported by several groups (20, 30, 44). In contrast, we observed an enrichment of *Lactobacillus* with early chronic SIVmac239 infection in RMs; the discrepancy between our findings and others may be due to the timing of our sampling during early chronic SIV infection. However, more interestingly, we also observed elevated fecal levels of *Lactobacillus* species in uninfected SMs relative to RMs. Together with our findings of low levels of Proteobacteria, this observation further supports a model in which SMs maintain a mucosal microbiome with lower propensity for inflammatory responses.

Within a research setting, we observed that increased levels of captivity of indoor-only standardized cage housing when compared to indoor-outdoor lifestyle reduced alpha diversity of GI microbiota, albeit modestly. Previous literature has reported a similar trend of reduced diversity between NHPs in captivity, or semi-captive lifestyles compared to those in the wild, although the changes were primarily attributed to dietary adaptations (27, 45). Our study shows that, within a diet-controlled setting, indoor housing is associated with a trend of reduced microbial diversity, overt bacterial community richness differences, and modest taxonomic disruptions within captive-born SMs. These data emphasize that intra-institutional or inter-institutional lifestyle differences should be controlled for in studies utilizing captive NHPs. However, we did not observe an effect of lifestyle on the microbial community structure between SIV-negative and SIV-positive SMs, and provided the rationale to pool SM data from the different lifestyle cohorts. The observation that housing had a greater impact than SIV infection status on the SM GI microbiome underscores the concept that the overall health of the gut environment is maintained during chronic lentiviral infection.

While reasonably powered, our study did have some drawbacks. On a technical level, serial longitudinal sampling at multiple time points has been used in macaque/SIV studies to control for the accelerated SIV pathogenesis and rapidly changing mucosal environment during acute infection of macaques. We were unable to choose a longitudinal design as de novo SIV infection of SMs is no longer allowable under current United States Fish and Wildlife Service regulations. Therefore, we chose a cross-sectional design, and obtained RM samples during the early chronic phase of infection, to avoid temporal fluctuations seen during the acute phase, and prior to long-term chronic infection, when gut health can be compromised. Other potential limitations of this study are: (i) the age differences between SMs and RMs; (ii) the differences in the duration of SIV infection; and (iii) the use of SMs from two locations with differing lifestyles (indoor and outdoor-indoor). The discrepancies in age were unavoidable, as colony breeding limitations, and the availability of SIV-infected SMs, necessitated the use of older animals, which are unavailable in the RM colony. As described above, the limited number of SIV-negative SMs maintained in an indoor lifestyle required the inclusion of outdoor-indoor SMs. We observed that there was a relatively modest variation, and an overall high level of concordance, in the bacterial community membership between SMs with differing environmental backgrounds, ages, and durations of SIV infection; suggesting that these variables had minimal influence on microbiome variability either prior to, or after SIV infection.

In this controlled diet-matched study, we report that the mucosal microbiome of SMs is comprised of enriched levels of beneficial anti-inflammatory gastrointestinal bacteria, and lower levels of inflammatory Proteobacteria, relative to non-natural host macaques. This difference in microbiota may contribute to the resistance developing the chronic immune activation which is seen with pathogenic infections. This study is the first to investigate the mucosal microbiome of the sooty mangabey species during SIV infection. We also found that the bacterial community structure in these species is highly stable after infection, likely reflecting the general health of the intestinal immune system and maintenance of the gut barrier. In HIV infection, a reproducible microbial dysbiosis is seen with alpha diversity loss, beta diversity expansion, and trends of specific taxa perturbances. This in turn, could be from mucosal integrity disruption or mucosal immunological-microbiota interactions which may lead to transient microbial community disruptions. Our findings, which show lower levels of Proteobacteria in natural hosts, provide support to the potential detrimental effect that pathobiont microbiota communities in potentiating a chronic inflammatory state during SIV infection. Previous literature have reported that prebiotics or probiotics with ART administration can enhance luminal immunological function and integrity which could potentially allow for avoidance of AIDS progression (42, 46); our data further supports a model in which lowering of pathobiont levels may reduce mucosal inflammation and immune dysfunction, and repair of gut barrier function.

## Materials and Methods

### Animals and Selection/Exclusion Criteria

Samples were collected from a total of 80 monkeys in accordance to institutional policies, which abide by the USDA Animal Welfare Regulations(47), the *Guide for the Care and Use of Laboratory Animals*(48), and the United States Fish and Wildlife Services Regulations at the Yerkes National Primate Research Center (YNPRC), an AAALAC-accredited institution. Fecal samples from fifty SMs were collected, including thirty SMs housed in indoor-outdoor enclosures at the YNPRC Field Station (Lawrenceville, GA), and twenty SMs housed indoor-only at YNPRC Main Center (Atlanta, GA) (**Table 1**). SMs at the YNPRC Field Station had equal groups of eight naturally-infected SIV positive and negative females, as well as seven naturally-infected SIV positive and negative males. The SM study cohort at the Main Center consisted of five SIV negative animals (2 females/3 males), and a total of fifteen SIVsmm positive SMs. Of the fifteen SIVsmm positive SMs at the Main Center (4 females/11 males) fourteen acquired SIVsmm through experimental intravenous infection (4 females/10 males) prior to November 2006 on previously approved IACUC protocols, and one (1 male) acquired SIVsmm naturally. Additionally, ten of the fifteen SIVsmm positive SMs at the Main Center, were previously given antiretroviral treatment (ART) which included a combination of tenofovir, emtricitabine, raltegravir, and darunavir on a previous study (26), however, all animals were more than 2.5 years post ART administration. Forty-eight SMs did not carry the homozygous CCR5-null alleles and two SMs had a CCR5 unknown genotype, and all fifty SMs were born at the YNPRC. All thirty SIVsmm-infected SMs were in a chronic phase of infection (>8 years except one naturally infected >3 years). Samples from thirty indoor-only housed RMs ranging from three to six years of age were collected at the YNPRC Main Center. RMs included equal groups of eight SIV positive and eight negative females as well as seven positive and seven negative males. All SIV positive RMs were experimentally infected with SIVmac239 (8 intrarectally/7 intravenously) on other IACUC approved protocols. RMs were in the early chronic phase of SIVmac239 infection (38-51 days) and naive to ART. Twenty-five RMs were born at YNPRC, five were born at either the Caribbean Primate Research Center, Mannheimer Foundation, or New England Primate Research Center and lived at YNPRC for a minimum of 12 months. All SMs or RMs that lived at the indoor facility have inhabited that location for a minimum of 5.5 months. All enrolled animals had not received antibiotics within 6 months, and 77 animals had not received antibiotics for over 12 months prior to sample collection **(Supplementary Table 1)** (49). To control for diet, all enrolled animals were uniformly fed LabDiet 5037 or 5038 (PMI Nutrition International, ST Louis, MO), which are identical formulated diets, that only vary by size. Additionally, nutritional enrichment are provided once daily and includes institutional approved vegetables, grains, or fruit to induce species-specific foraging behaviors. NHPs are provided with manipulable objects (i.e. Kong toys®, Nylabones®, PVC objects, metal triangles, wooden blocks or mirrors) for enrichment. Indoor-outdoor enclosures have physical enrichment including but not limited to barrels, swings, fire hose, climbing structures, milk crates, and perches. Singly housed animals are given additional destructible enrichment (i.e. paper or cardboard) and foraging-boards to enhance species-specific behaviors.

### Collection of NHP fecal samples

Approximately 1 gram of feces were collected opportunistically from all animals within one hour of defecation on grossly uncontaminated surfaces (areas with no other pre-existing urine, water, food, natural substrates, or feces). Sample collection was performed in a manner in which fecal surfaces in contact with grossly uncontaminated surfaces were avoided. The fecal samples were transferred into DNA-free 15 ml conical tubes at the time of collection and stored for future processing at −80 °Celsius.

### Genomic DNA Extraction and 16S rRNA NGS library construction

Total DNA was extracted from frozen stool samples using the E.Z.N.A. Stool kit (Omega Bio-tek, GA). Libraries were prepared using the Bioo Scientific NextFlex 16s V4 Amplicon-Seq kit (Bioo Scientific Corporation, TX). A first PCR amplification was performed using customized primers that target the V4 domain and a second PCR to integrate relevant flow cell binding domains and unique sample indices. The libraries generated were validated using a DNA tape on the Agilent 4200 TapeStation and quantified using fluorescence based method. The libraries were normalized and pooled. Clustering and sequencing of the libraries occurred on an Illumina miSeq employing a 301PE run.

### Bioinformatic and Statistical Analyses

Per sample “.fastq” read files were demultiplexed in the Illumina BaseSpace Cloud environment and downloaded for processing and analysis. Reads were filtered and processed using QIIME 2 (57) within a conda environment under Linux version qiime2-2019.4. using the dada2 [doi://10.1038/nmeth.3869] plugin “denoise-paired” method producing a binned feature table of representative sequences. Samples with feature counts of more than 10000 were retained for downstream analysis with only one sample removed (RMo16) with a feature count of 2135 was excluded from the alpha and beta diversity measurement plots but was represented in all other figures. Sequences were classified using the Greengenes_13_8 99% database.

Taxonomic relative abundances are calculated relative to all features classified as bacterial. Alpha diversity was calculated for Shannon and Chao1 indices. Beta diversity was evaluated using unweighted UniFrac distances for intra-host species viral status comparisons and with the Bray-Curtis distance for cross host species and environmental comparisons. Significance of alpha diversity measure and taxa relative abundance differences was performed in R using the core two-sample Wilcoxon Rank Sum test.

## Acknowledgements

We would like to acknowledge veterinary and colony management staff: Tracy Meeker, Amanda Cerqueda and Madison Prestipino; Bhavani Madeti for generation of 16S rRNA libraries; and Diane Carnathan for animal records. This study was funded in part by ORIP/OD P51OD011132 to the YNPRC.

---

**Figure S1. Comparison of the fecal microbiome in indoor versus outdoor-indoor SMs shows a reduction in biodiversity and mild phylogenetic taxonomic disruption.**

**A)** Bar graph representation of phylogenetic phyla (top figure) and family (bottom figure) bacterial community acquisition. Each individual SM represents a single bar and are grouped according to housing; indoor-outdoor on left, indoor-only on right. **B**) Chao1 index (top figure) a total OTU richness measurement for alpha diversity comparing indoor-only with indoor-outdoor housing. Shannon diversity (bottom figure) an alpha diversity metric, measuring bacterial community richness and evenness between cohorts, this representation compares indoor-outdoor (red) with indoor only (black) housing. Both alpha diversity metrices were compared using the paired Wilcoxon rank sum test. **C)** Relative abundance of the predominant phyla Firmicutes, Bacteroidetes, Spirochaetes, Proteobacteria represented by box plots comparing indoor-outdoor (red) and indoor-only (black) SM cohorts compared by the paired Wilcoxon rank sum test. **D)** Relative abundance of genera *Prevotella, Lactobacillus*, and *Streptococcus* represented by box plots comparing indoor-outdoor (red) and indoor-only (black) SM cohorts compared by the paired Wilcoxon rank sum test. **E)** Principal coordinate plot of Bray-Curtis a dissimilarities measurement of beta diversity comparing indoor-outdoor (red) and indoor-only (black) SM cohorts.

